# *Dirofilaria immitis* and *D. repens* in Europe: a systematic literature review on vectors, host range, and the spatial distribution in the 20th and 21st century

**DOI:** 10.1101/2025.02.17.638693

**Authors:** Carolin Hattendorf, Renke Lühken

## Abstract

**Background:** *Dirofilaria immitis* and *D. repens* are mosquito-borne nematodes with dogs as primary hosts, but other mammalian species including humans can be also infected. In the last century, circulation of both pathogens was predominantly restricted to Southern Europe. However, different studies indicated a potential establishment in Central, Eastern and Western parts of Europe as an increasing threat to animal and human health.

**Methods:** We conducted a systematic literature review of publications reporting *D. immitis* and *D. repens* screening in mosquitoes and mammalian vertebrates in Europe. These data were used to analyse the range of vectors and hosts and for a comparison of the spatial distribution between the 20^th^ and 21^st^ century.

**Results:** Both nematodes appear to have a high overlap of *Aedes*, *Anopheles* and *Culex* vector species, which are abundant in Europe. Most *D. immitis* infections were reported in dogs, while *D. repens* predominated in humans. *Dirofilaria immitis* infections were detected in a wider range of wild and zoo animals. Compared to the last century, many more countries especially in Central Europe were affected by *Dirofilaria* spp. circulation, illustrating a significant spread over the last 20 years.

**Conclusion:** Our findings suggest that *D. immitis* and *D. repens* are a growing health concern for animals and humans in Europe. Continuous globalisation and climate warming will probably lead to a further spread and increased circulation in the future. All data are made available open access, which will enable further analysis in the future.

## Introduction

Two *Dirofilaria* species are present in Europe: *D. immitis* and *D. repens* (1). Both circulate in an enzootic cycle between mosquitoes and domestic dogs, although other carnivores like Red Foxes and Grey Wolves can also be infected (e.g. (2–5)). Mosquitoes are infected with microfilaria during blood-feeding on an infected host, which then develop to infective larvae in susceptible vectors (6). *Dirofilaria* can be transmitted to other mammals, such as humans and rodents, although these are generally ‘dead-end’ hosts (6), i.e. no development of microfilaria occurs. *Dirofilaria immitis* localise in the pulmonary arteries of dogs, where they sexually reproduce and release microfilariae into the bloodstream (1,7). Infections can lead to severe disease in dogs and cats with symptoms ranging from chronic cough to heart failure (8,9). In humans, *D. immitis* mostly forms pulmonary nodes, which are generally asymptomatic, but frequently mistaken with lung cancer in radiography (6). However, some humans develop severe symptoms including fever, chest pain, coughing, haemoptysis, wheezing arthralgia or malaise (10). *Dirofilaria repens* infections generally localises subcutaneously (1,6). Approximately 35 % of human *D. repens* infections occur in the ocular region, which can lead to impaired or a complete loss of vision (11). Around 10 % of affected patients suffer permanent complications like retinal detachment or glaucoma (12). Notably, there have been a few reported cases where viable *D. repens* microfilariae have been found in the blood stream of infected humans (13–17), but these seem to be rare exceptions. The majority of human *Dirofilaria* infections in Europe are caused by *D. repens* (18), while the majority of reported *Dirofilaria* cases in dogs are *D. immitis* (1). However, it has to be noted that *D. immitis* is easier to diagnose in dogs because it more often leads to severe symptoms in dogs and respective tests are available (19).

First cases of human dirofilariosis presumably were diagnosed in 1566 in a Portuguese girl (20) and 1626 in an Italian dog (21) for *D. repens* and *D. immitis*, respectively. In the 20th century, autochthonous circulation of these parasites was predominantly reported from the Southern parts of Europe, but currently there are increasing reports of a spread towards Central, West and East Europe (22). Many previously *Dirofilaria*-free countries are now considered endemic (23). Climate warming is thought to be the main reason, allowing the successful development of the nematodes in the mosquito (24–26). Another important factor is the movement of dogs in Europe, which was made considerably easier with European regulations for traveling with pets (27). To gain a better picture of the vector range and spatial expansion of *D. immitis* and *D. repens* in Europe over the last two centuries, we conducted a systematic literature review of *Dirofilaria* data in mosquitoes and vertebrate hosts, including the collection of different metadata (e.g. sampling time and site).

## Methods

All published articles matching the keyword ‘dirofilaria’ in any search field recorded in PubMed (28) were extracted on 24.01.2022. Papers were selected using the following inclusion criteria: 1) article language English or German, 2) a host was diagnosed with an acute infection of *Dirofilaria* spp., i.e. excluding studies only screening antibodies, and 3) the sampling was conducted in Europe. The following information was extracted from each publication: country, date of diagnosis/sampling, sampling location, host species, travel history, screening method, number of tested and number of positive specimens per *Dirofilaria* species. In addition, for mosquito studies the mosquito trap and pooling information (pool size, body part, etc.) were noted.

If the date of diagnosis was not specified, the date of publication was used and if only a sampling period was given, the total number of cases was split evenly across the sampling years. The accuracy of the sampling locations was classified to decide which level of the Nomenclature of Territorial Units for Statistics (NUTS) classification of the European Union (29) was used for visualisation of parasite distribution in humans, dogs and other vertebrate hosts: ‘very high’ (coordinates or address, NUTS-3 level), ‘high’ (town or specific area, NUTS-2 level), ‘medium’ (hospital or greater area (e.g. county), NUTS-1 level), and ‘low’ (country, NUTS-0 level). For the spatial analysis of the *Dirofilaria* distribution, we only included reports with unremarkable travel history. However, many studies did not include any information on the travel history. Therefore, we also conducted the spatial visualisation with all unremarkable and unknown travel history for the supplement. Reports with a known travel history were excluded from analysis. Furthermore, we compiled visual summaries of country-specific *Dirofilaria* screening results from mosquitoes and less common vertebrate hosts, excluding humans and dogs. All computational analysis was performed in R (Version: 4.2.2) using the R-Studio IDE (Version:2022.12.0) (30). Additionally, functions from the following packages were used for data preparation, visualization and analysis: terra (31), tidyterra (32), geodata (33) readxl (34), ggpubr (35), plyr (36), dplyr (37), and ggplot2 (38).

## Results

A total of 3,847 publications were extracted from PubMed. Of these, 473 (12.3 %) matched our inclusion criteria. We observed an increase in publications reporting *Dirofilaria* from the beginning of the 1990s and another increase in the mid-2000s (Fig. 1).

**Fig. 1:**
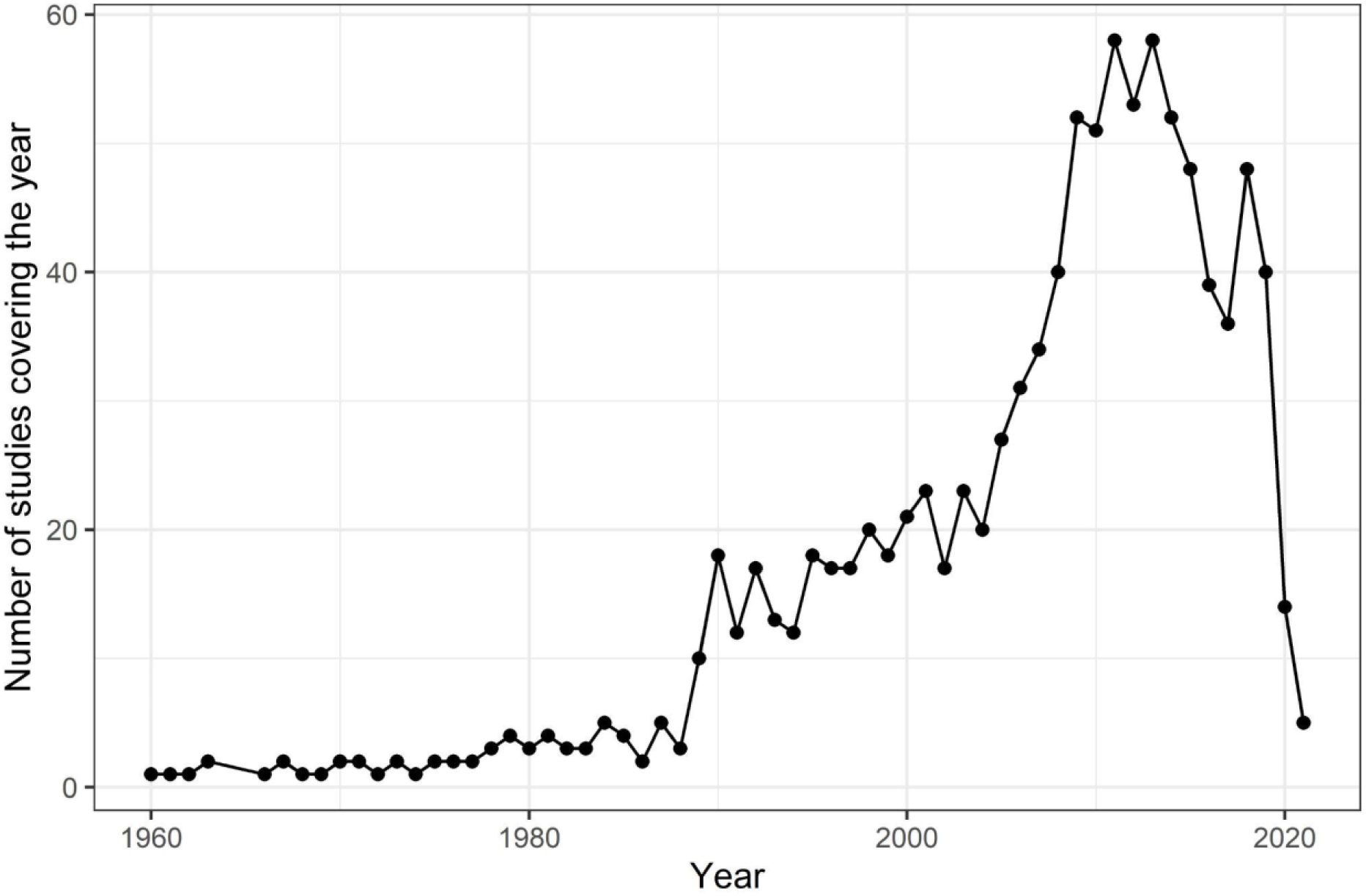
Number of studies reporting *Dirofilaria immitis* and *D. repens* in Europe.

38 publications (8.0 %) included screenings of mosquitoes for *Dirofilaria* with a total of 1,658,041 specimens tested over 62 mosquito taxa collected in 14 different European countries (Fig. 2). *Dirofilaria immitis* was detected in 17 different mosquito taxa from 12 countries, most frequently in *Culex pipiens* s.l. (11 countries) and *Aedes caspius* (7 countries). *Dirofilaria repens* infections were reported for 31 different mosquito species from 13 countries with *Aedes vexans* (8 countries), *Cx. pipiens* s.l. (6 countries) and *Anopheles maculipennis* s.l. (6 countries) most frequently found positive. A total of 15 mosquito taxa were found positive for both *Dirofilaria* species. *Dirofilaria immitis* was exclusively detected in *Ae. behningi*, while *D. repens* was exclusively found in 16 different taxa of the *Aedes*, *Anopheles*, *Culiseta* and the *Uranotaenia* genus, e.g. *Ae. cantans*, *An. claviger*, *Cs. annulata* or *Ur. unguiculata*. Most studies on *Dirofilaria* prevalence in mosquitoes focused on Southern and Eastern Europe, but some studies also confirmed autochthonous circulation in Central Europe, e.g. Austria or Germany.

**Fig. 2:**
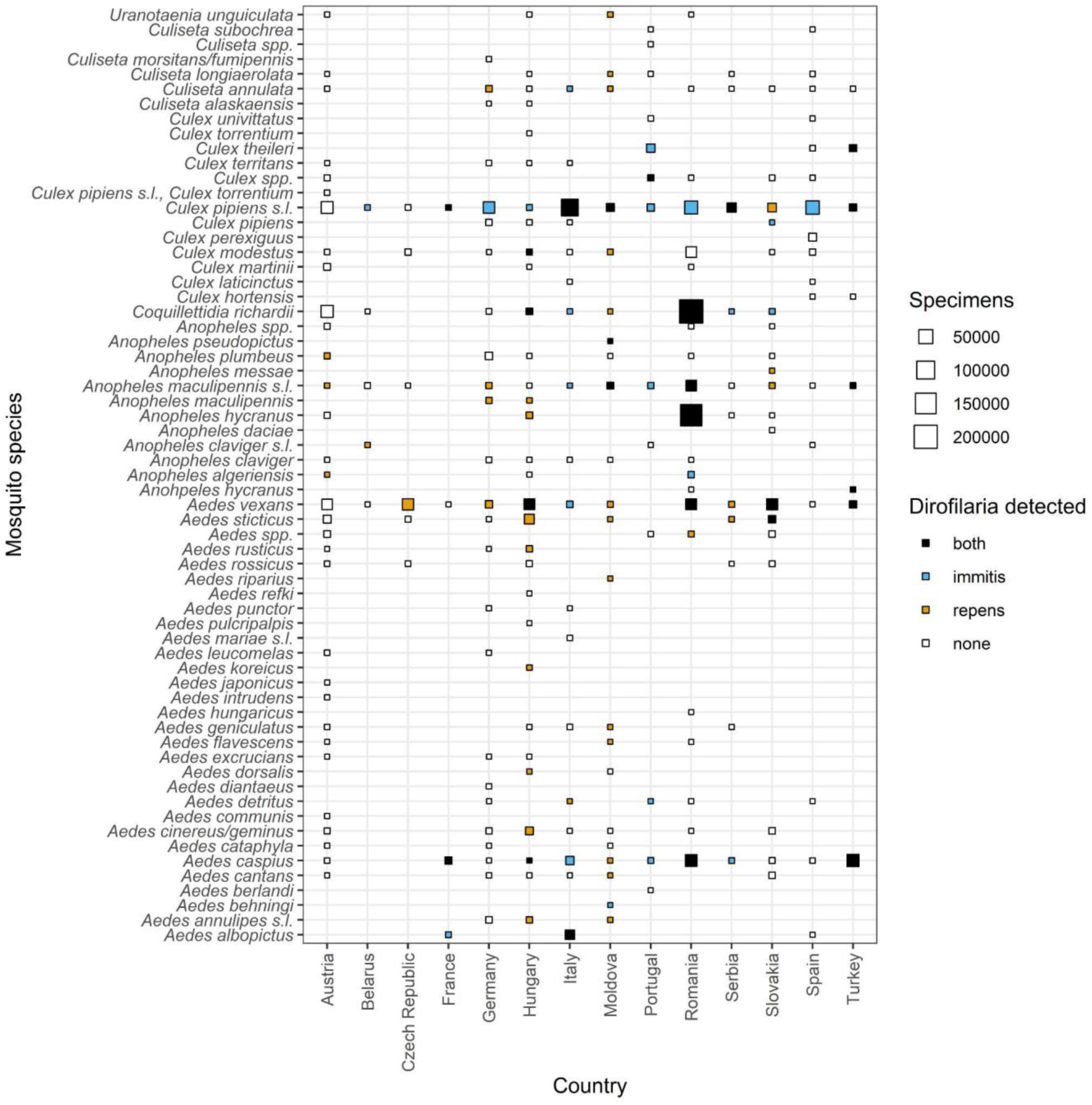
*Dirofilaria immitis* and *D. repens* reports in mosquitoes for different European countries.

198 publications (41.9 % of included publications) reported dog infections with a total of 11,713 cases. Of these, 7,757 (66.2 %) were identified as *D. immitis*, 3,948 (33.7 %) as *D. repens*, and eight (0.1 %) were not further differentiated *Dirofilaria* species. In 199 publications (42.1 %), human *Dirofilaria* spp. infections were described, summing up to 2,555 reported human cases, of which the majority of 2,438 (95.4 %) was *D. repens*, followed by 95 (3.7 %) not further specified *Dirofilaria* spp. and 22 (0.9 %) *D. immitis*. Only 33 publications (7.0 %) reported *Dirofilaria* infections in cats (278 cases): 252 (90.1 %) *D. immitis*, 24 (8.6 %) *D. repens* and two (0.7 %) not further specified *Dirofilaria* species. In addition, 59 publications (12.5 %) described *Dirofilaria* infection in other mammals, the majority of which were caused by *D. immitis* (Fig. 3). These studies predominantly focused on domestic cats (34 publications, 7.2 %) and Red Foxes (15 publications, 3.2 %). In addition, *Dirofilaria* were detected in a wide variety of wild carnivores (e.g. Golden Jackal, Grey Wolf or Eurasian Otter) and zoo animals (e.g. Lion or California Sea Lion). A wider variety of vertebrate hosts was studied in Slovakia, Serbia and Romania, while studies in other countries focused on few potentially infected species like Red Foxes or only reported single cases.

**Fig. 3:**
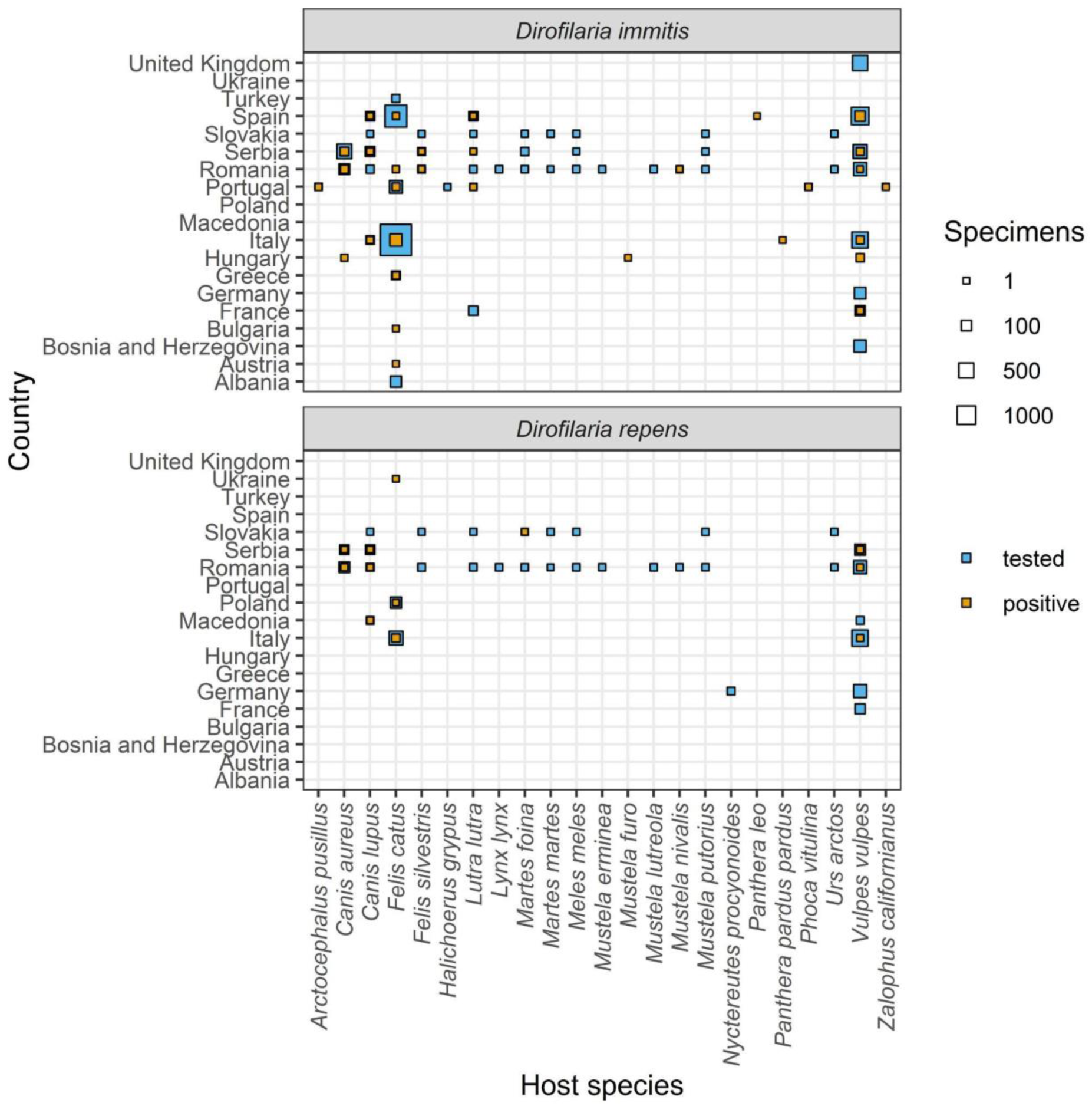
*Dirofilaria immitis* and *D. repens* reports in vertebrates except humans and dogs for different European countries.

**Fig. 4:**
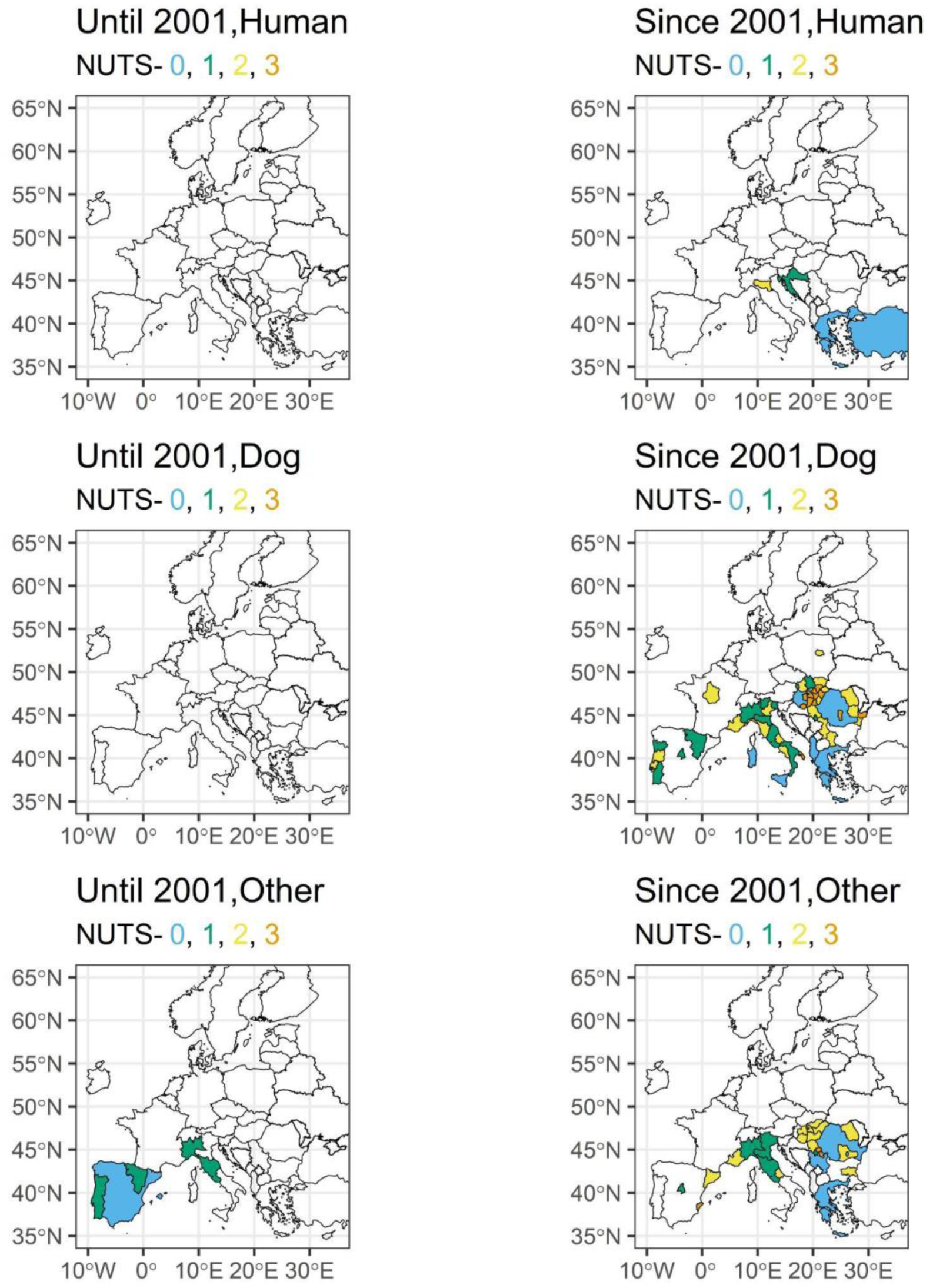
*Dirofilaria immitis* cases in humans, dogs and other mammals with unremarkable travel history in Europe until and since 2001 at different geographical levels.

**Fig. 5:**
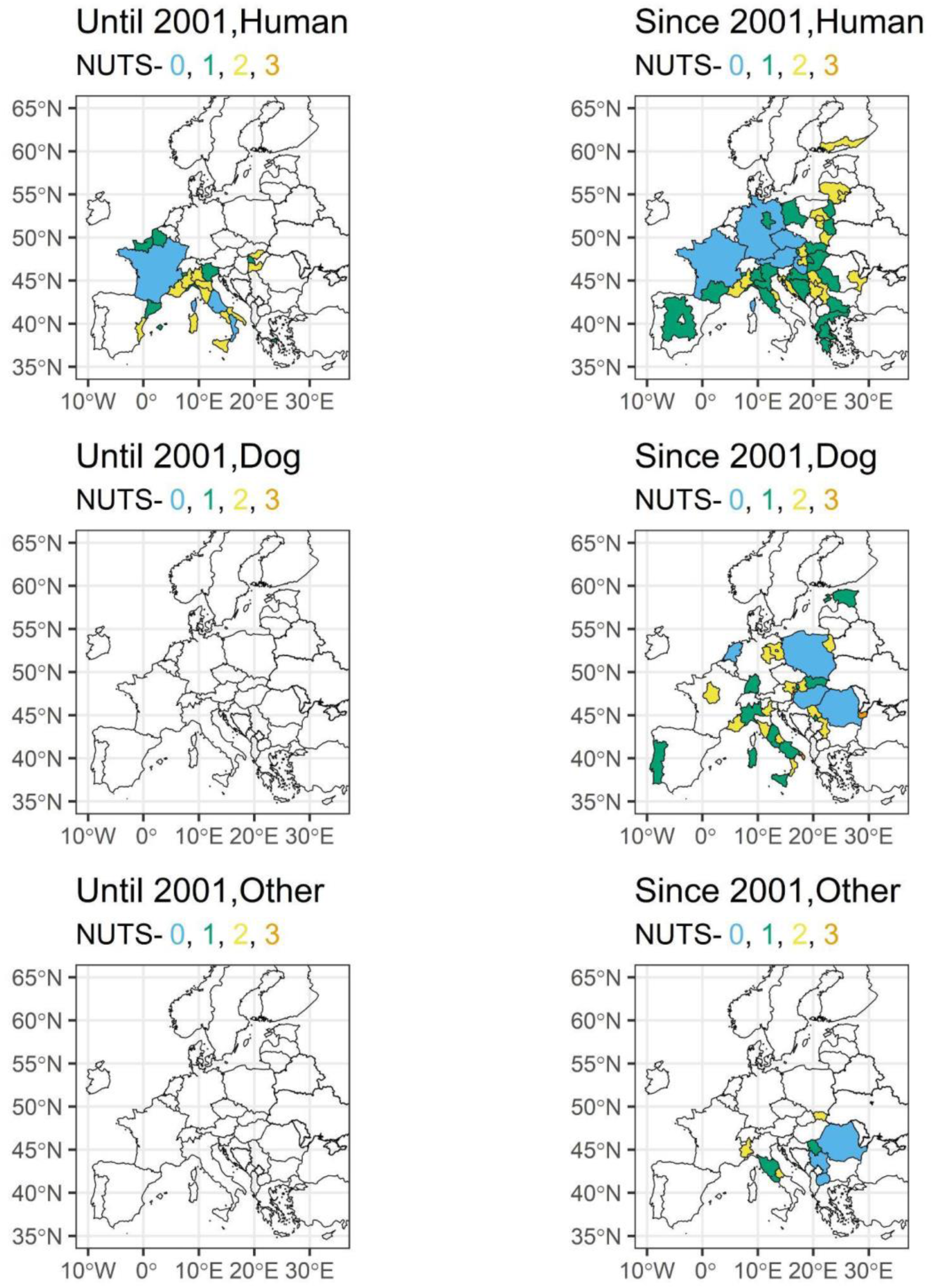
*Dirofilaria repens* cases in humans, dogs and other mammals with unremarkable travel history in Europe until and since 2001 at different geographical levels.

Only focusing on the studies with unremarkable travel history, the majority of the few *D. immitis* cases in dogs and other mammals until 2001 were recorded in Southern Europe, particularly in Spain, Italy and Portugal (Fig. 2; see supplementary file 1 and supplementary file 2 for visualisation of all cases with unremarkable and unknown travel history). No human cases were reported before 2001. In the 21^st^ century, *D. immitis* infections were found in most countries of South and Central Europe and even in Central Europe, such as Poland and France. A wide distribution in particular was confirmed in dogs and other mammals for wide parts of Eastern Europe and Italy. *Dirofilaria repens* infections, especially looking into human cases, were reported much more widespread than *D. immitis* already during the 20^th^ century in particular for various regions in Italy and France, while dogs were only tested positive in Italy and Spain (Fig. 3). We observed a strong increase of affected countries for both, humans and dogs, including countries in Eastern and Southern Europe (e.g. Ukraine, Slovakia, Greece), but also Central Europe including the Netherlands, Germany or Poland. The most Northern infection was reported in humans from Finland.

## Discussion

The number of publications reporting *Dirofilaria* spp. infections have increased in the last two decades compared to the previous century (18,22). This is most likely driven by both, increased research and awareness, but also the spread of the parasites (25,39). *Dirofilaria immitis* and *D. repens* have to be considered endemic in countries that were considered to be *Dirofilaria*-free in the 20^th^ century, e.g. Czech Republic (40,41). The spread of competent vector species probably does not play a major role here. There is a huge overlap between the vector species for *D. immitis* and *D. repens*, which are widespread in Europe and show host-feeding patterns with a substantial proportion of mammals, e.g. *Cx. pipiens* s.l. or *An. maculipennis* s.l. (42–44). Interestingly, the exotic *Ae. albopictus* was much more often reported to be infected with *D. immitis* than *D. repens*. This mosquito species has been implicated as an important driver of the spread of *Dirofilaria* (22,26,45).

Most infections were reported from dogs as the primary host of *Dirofilaria* (1). The majority of these cases were caused by *D. immitis*, which is well known to cause a more severe disease in dogs compared to *D. repens*, leading to a higher probability of diagnosis (7). Additionally, rapid tests are only available for *D. immitis* and not *D. repens.* Therefore, *D. repens* might be underreported and its actual prevalence among dogs is probably higher (11). In contrast, the overwhelming majority of cases in humans were caused by *D. repens*, confirming previous observations that most human *Dirofilaria* infections in Europe are caused by this species (46,47). The reason for this remains unclear, given that human infections with *D. immitis* are regularly reported, particularly in North America (48). One hypothesis suggested that European *D. immitis* might be genetically distinct from *D. immitis* found in other regions, making it less capable of surviving within humans (11). However, this hypothesis has later been disproven (49,50). Another explanation could be that *D. repens* influences the circulation of *D. immitis*, e.g. it has been shown for Southern Italy that *D. repens* impedes the spread of *D. immitis* in dogs (51). If this plays a general epidemiological role and if this is also true for humans requires further research. Furthermore, it has been proposed that *D. repens* is more difficult to control, because, as mentioned above, rapid tests are only available for *D. immitis* and current preventative and curative treatments are designed for *D. immitis* and are not as effective against *D. repens* (18). Additionally, *D. repens* infections are often asymptomatic in dogs which might lead to a longer time period where a dog is infective, and a mosquito can ingest and transmit the parasite to further hosts (6,52).

Besides dogs and humans, there were also several reports of infections in cats, although it is assumed that cats do not play an important role for *Dirofilaria* transmission (53). Similarly, several other mammalian species diagnosed with an infection were held in zoos or as pets, allowing diagnosis (54–57). Furthermore, there are some wild animals in which *Dirofilaria* infections were identified, predominantly in canids like Red Foxes (4,5,58–69), Golden Jackals (59–62,70,71), and Grey Wolves (2,3,58,59,62,72,73). Zoo and wild animals were almost always infected by *D. immitis*, which again might be because *D. immitis* in comparison to *D. repen*s infections more often leads to severe symptoms, corresponding test kits are available or because *D. immitis* has a broader host range.

It is undeniable that both, *D. immitis* and *D. repens*, are spreading in Europe and more humans and animals are at risk of infection. In part this might be also a diagnostic artefact, i.e. imported and travelling dogs are more routinely tested, which leads to more detection (39). Transport of pets has significantly increased during the 21^st^ century as a consequence of the Pet Travel Scheme, which was introduced by the EU in 2000 and made travel of companion animals significantly easier and led to an increase in imported cases (27). Another reason is the continuously high number of stray dogs in some countries, which are not subject to regular treatment and act as reservoirs for the parasites, e.g. countries with many stray dogs, such as Romania or Bulgaria continue to regularly report *Dirofilaria* spp. cases (74–76). However, probably one of the most important factors for the spread of *Dirofilaria* is climate warming. Higher temperatures lead to faster development of *Dirofilaria* larvae inside the mosquito vector (77,78). Prolonged warm periods extend the transmission season (26). There is a significant increase in areas at risk, especially in more Northern countries. This spread has been predicted since the early 2000s (25,52) and with continuous climate warming will further increase in the future. Finally, increasing temperatures in Europe also allowed the widespread establishment of exotic mosquito species such as *Ae. albopictus* (79–81), which is an important vector for *D. immitis* and *D. repens* and the establishment of the vector species in numerous European regions has been linked to increased *Dirofilaria* circulation (22).

## Conclusion

*Dirofilaria immitis* and *D. repens* are an increasing threat to veterinary and public health in Europe. Both parasites have dramatically expanded their circulation area and are now endemic in areas that were considered *Dirofilaria-*free only one or two decades ago (23). The warming climate and the abundant presence of competent vectors allows the establishment of the parasites in Central Europe, e.g. Germany and Poland. Due to their rising relevance in animal and human health, a Europe-wide unified surveillance system similar to the system in the United States (82) should be implemented in order to better understand the change of circulation patterns and to plan and execute preventative strategies, e.g. dog treatment. All data and code are provided as open access, allowing for future analyses.

## Ethics approval and consent to participate

Not applicable.

## Consent for publication

Not applicable.

## Availability of data and materials

The datasets supporting the conclusions of this article and all codes used for data analysis are available at https://github.com/luehkenecology/dirofilaria_review_europe

## Author contributions

Conceptualization: RL; data collection: CH; data analysis: CH, RL; writing: CH, RL; all authors read and approved the final manuscript.

## Acknowledgements

Not applicable.

---

**Supplementary file 1:**
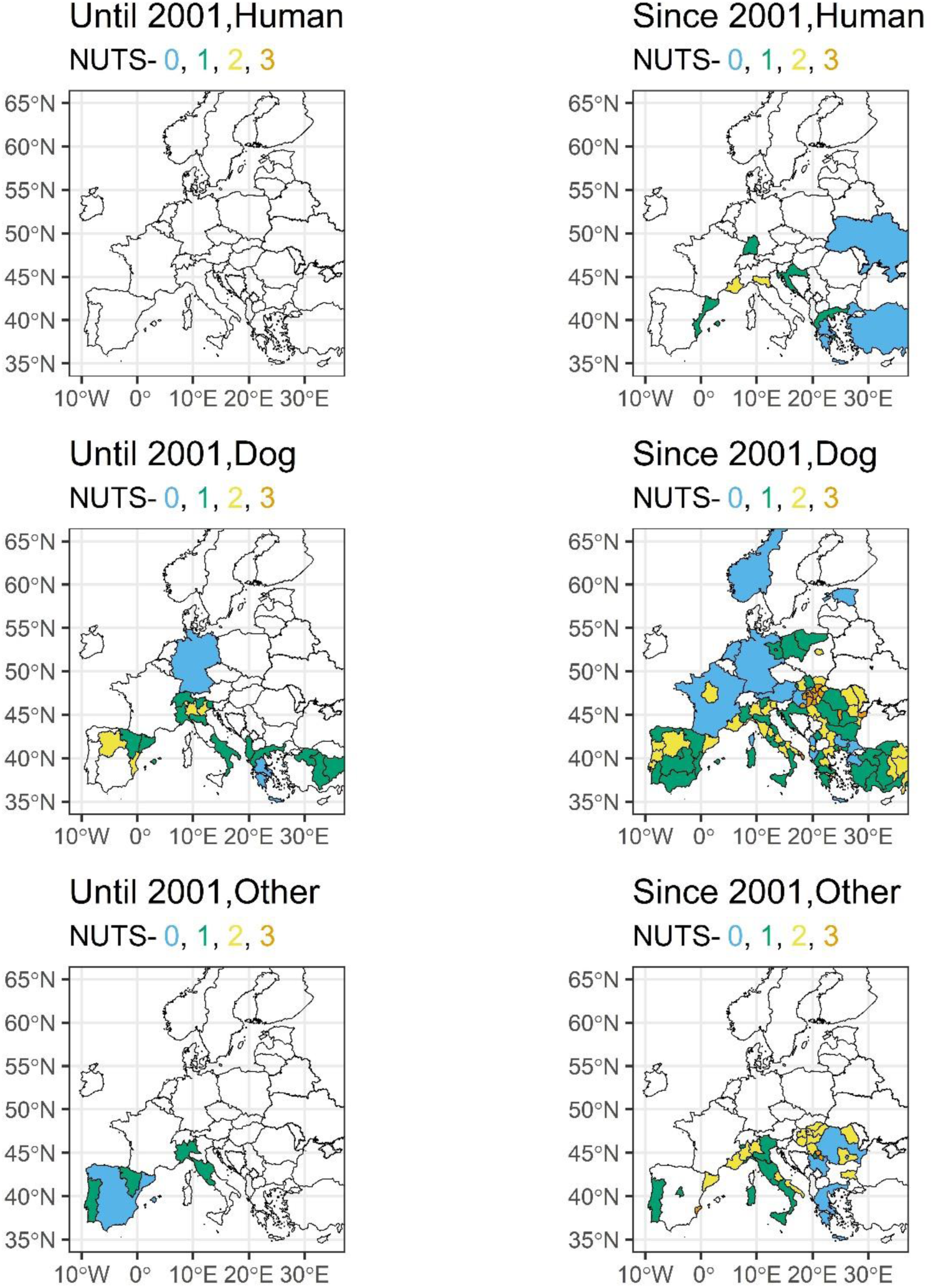
*Dirofilaria immitis* cases in humans, dogs and other mammals with unremarkable and unknown travel history in Europe until and since 2001 at different geographical levels.

**Supplementary file 2:**
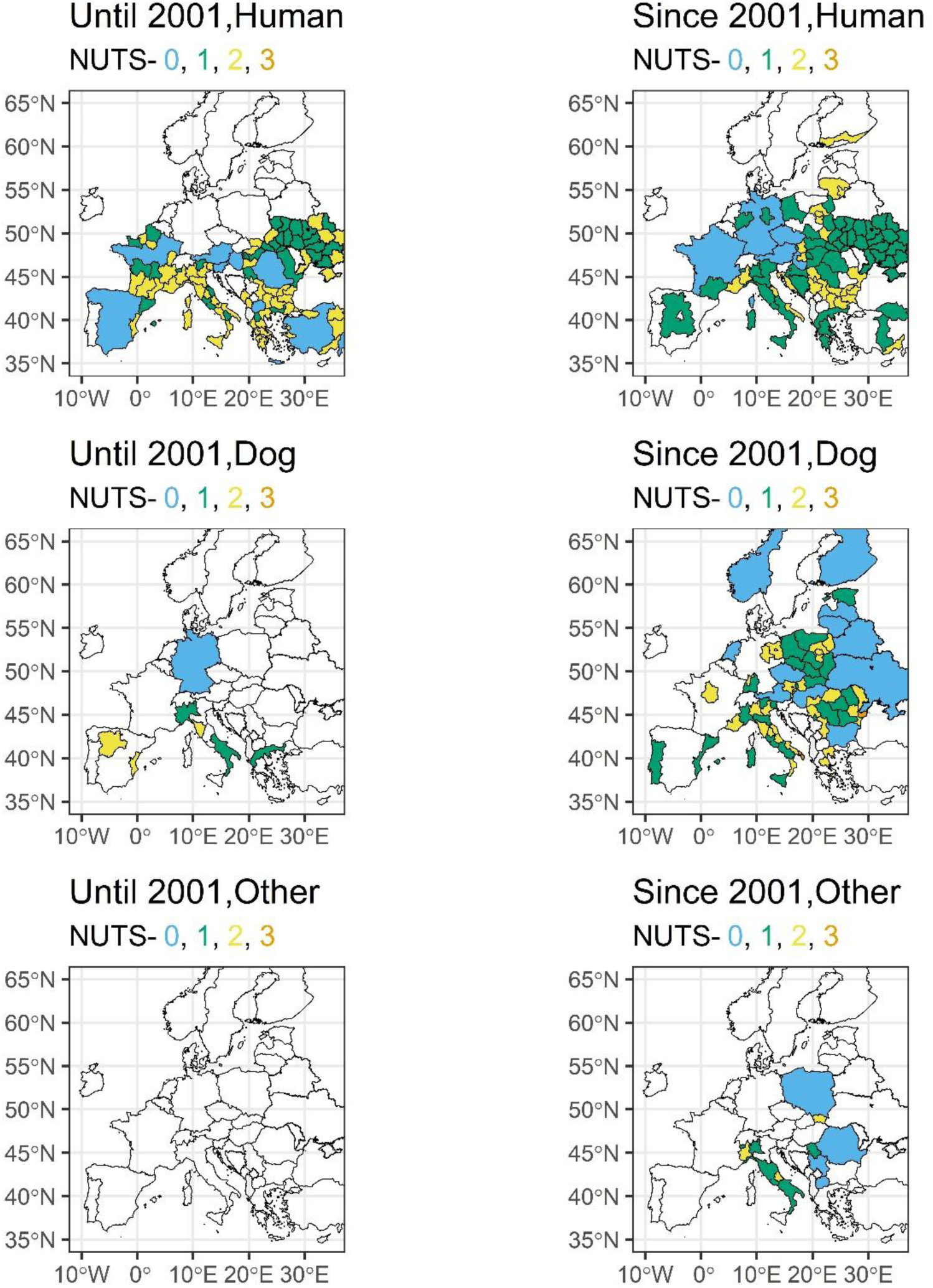
*Dirofilaria repens* cases in humans, dogs and other mammals with with unremarkable and unknown travel history in Europe until and since 2001 at different geographical levels.

## References

1. McCall JW, Genchi C, Kramer LH, Guerrero J, Venco L. Heartworm disease in animals and humans. Adv Parasitol. 2008;66:193–285.

2. Moroni B, Rossi L, Meneguz PG, Orusa R, Zoppi S, Robetto S, et al. Dirofilaria immitis in wolves recolonizing northern Italy: are wolves competent hosts? Parasit Vectors. 2020 Sep 22;13(1).

3. Segovia JM, Torres J, Miquel J, Llaneza L, Feliu C. Helminths in the wolf, Canis lupus, from north-western Spain. J Helminthol. 2001 Jun;75(2):183–92.

4. Medkour H, Laidoudi Y, Marié J Lou, Fenollar F, Davoust B, Mediannikov O. Molecular investigation of vector-borne pathogens in red foxes (Vulpes vulpes) from southern France. J Wildl Dis. 2020;56(4):837–50.

5. Liesner JM, Krücken J, Schaper R, Pachnicke S, Kohn B, Müller E, et al. Vector-borne pathogens in dogs and red foxes from the federal state of Brandenburg, Germany. Vet Parasitol. 2016;224:44–51.

6. Simón F, Siles-Lucas M, Morchón R, González-Miguel J, Mellado I, Carretón E, et al. Human and animal dirofilariasis: the emergence of a zoonotic mosaic. Clin Microbiol Rev [Internet]. 2012 Jul [cited 2023 Jun 26];25(3):507–44. Available from: https://pubmed.ncbi.nlm.nih.gov/22763636/

7. Bowman DD, Atkins CE. Heartworm biology, treatment, and control. Vet Clin North Am Small Anim Pract. 2009 Nov;39(6):1127–58.

8. Garrity S, Lee-Fowler T, Reinero C. Feline asthma and heartworm disease: Clinical features, diagnostics and therapeutics. J Feline Med Surg. 2019 Sep 1;21(9):825–34.

9. Ames MK, Atkins CE. Treatment of dogs with severe heartworm disease. Vet Parasitol. 2020 Jul 1;283:109131.

10. Saha BK, Bonnier A, Chong WH, Chieng H, Austin A, Hu K, et al. Human Pulmonary Dirofilariasis: A Review for the Clinicians. Am J Med Sci. 2022 Jan 1;363(1):11–7.

11. Genchi C, Kramer LH, Rivasi F. Dirofilarial infections in Europe. In: Vector-Borne and Zoonotic Diseases. 2011. p. 1307–17.

12. Avdiukhina TI, Lysenko AI, Supriaga VG, Postnova VF. Dirofilariasis of the vision organ: registry and analysis of 50 cases in the Russian Federation and in countries of the United Independent States. Vestn Oftalmol. 1996;112(3):35–9.

13. Negahban S, Daneshbod Y, Atefi S, Daneshbod K, Sadjjadi SM, Hosseini SV, et al. Dirofilaria repens diagnosed by the presence of microfilariae in fine needle aspirates: a case report. Acta Cytol. 2007;51(4):567–70.

14. Potters I, Vanfraechem G, Bottieau E. Dirofilaria repens Nematode Infection with Microfilaremia in Traveler Returning to Belgium from Senegal. Emerg Infect Dis. 2018 Sep 1;24(9):1761.

15. Petrocheilou V, Theodorakis M, Williams J, Prifti H, Georgilis K, Apostolopoulou I, et al. Microfilaremia from a Dirofilaria-like parasite in Greece. Case report. APMIS. 1998 Feb;106(2):315–8.

16. Nozais JP, Bain O, Gentilini M. [A case of subcutaneous dirofilaria (Nochtiella) repens with microfilaremia originating in Corsica]. Bull Soc Pathol Exot. 1994;87(3):183–5.

17. Pupić-Bakrač A, Pupić-Bakrač J, Beck A, Jurković D, Polkinghorne A, Beck R. Dirofilaria repens microfilaremia in humans: Case description and literature review. One Health. 2021 Dec 1;13:2352–7714.

18. Genchi C, Kramer L. Subcutaneous dirofilariosis (Dirofilaria repens): an infection spreading throughout the old world. Parasit Vectors. 2017 Nov 9;10(Suppl 2).

19. Capelli G, Genchi C, Baneth G, Bourdeau P, Brianti E, Cardoso L, et al. Recent advances on Dirofilaria repens in dogs and humans in Europe. Parasit Vectors. 2018;11(1):1–21.

20. Lusitano A. Curationum Medicinalium Centuria Septima. In: Venetiis: apud Vincentium Valgresium, curatio. 1566. p. 106.

21. Birago F. Trattato cinegetico, ovvero della caccia. In: Sfondrato V. Milan, Italy; 1626. p. 77.

22. Genchi C, Kramer LH. The prevalence of Dirofilaria immitis and D. repens in the Old World. Vet Parasitol. 2020;280:108995.

23. Masny A, Gołąb E, Cielecka D, Sałamatin R. Vector-borne helminths of dogs and humans - focus on central and eastern parts of Europe. Parasit Vectors. 2013 Feb 22;6(1):38.

24. Széll Z, Bacsadi Á, Szeredi L, Nemes C, Fézer B, Bakcsa E, et al. Rapid spread and emergence of heartworm resulting from climate and climate-driven ecological changes in Hungary. Vet Parasitol. 2020 Apr 1;280.

25. Genchi C, Rinaldi L, Cascone C, Mortarino M, Cringoli G. Is heartworm disease really spreading in Europe? Vet Parasitol. 2005 Oct 24;133(2–3):137–48.

26. Genchi C, Rinaldi L, Mortarino M, Genchi M, Cringoli G. Climate and Dirofilaria infection in Europe. Vet Parasitol. 2009 Aug 26;163(4):286–92.

27. Trotz-Williams LA, Trees AJ. Systematic review of the distribution of the major vector-borne parasitic infections in dogs and cats in Europe. Vet Rec. 2003 Jan 25;152(4):97– 105.

28. National Library of Medicine. PubMed [Internet]. [cited 2023 Jul 28]. Available from: https://pubmed.ncbi.nlm.nih.gov/

29. Background - NUTS - Nomenclature of territorial units for statistics - Eurostat [Internet]. [cited 2023 Jul 28]. Available from: https://ec.europa.eu/eurostat/web/nuts/background

30. R Core Team. R: A Language and Environment for Statistical Computing [Internet]. Vienna, Austria: R Foundation for Statistical Computing; 2023. Available from: https://www.R-project.org

31. Hijmans R. terra: Spatial Data Analysis_. R package version 1.7-78 [Internet]. 2024. Available from: https://CRAN.R-project.org/package=terra

32. Hernangómez D. Using the tidyverse with terra objects: the tidyterra package. J Open Source Softw. 2023 Nov 10;8(91):5751.

33. Hijmans RJ, Barbosa M, Ghosh A, Mandel A. geodata: Download Geographic Data. R package version 06-2 [Internet]. 2024; Available from: https://CRAN.R-project.org/package=geodata

34. Wickham H, Bryan J. readxl: Read Excel Files [Internet]. 2023. Available from: https://CRAN.R-project.org/package=readxl

35. Kassambara A. ggpubr: “ggplot2” Based Publication Ready Plots [Internet]. 2023. Available from: https://CRAN.R-project.org/package=ggpubr

36. Wickham H. The Split-Apply-Combine Strategy for Data Analysis. Journal of Statistical Software. 2011;40(1):1–29.

37. Wickham H, François R, Henry L, Müller K, Vaughan D. dplyr: A Grammar of Data Manipulation [Internet]. 2023. Available from: https://CRAN.R-project.org/package=dplyr

38. Wickham H. ggplot2: Elegant Graphics for Data Analysis. Springer-Verlag New York,; 2016.

39. Genchi C, Bowman D, Drake J. Canine heartworm disease (Dirofilaria immitis) in Western Europe: survey of veterinary awareness and perceptions. Parasit Vectors. 2014 Apr 29;7(1).

40. Morchón R, Montoya-Alonso JA, Rodríguez-Escolar I, Carretón E. What Has Happened to Heartworm Disease in Europe in the Last 10 Years? Pathogens. 2022 Sep 1;11(9).

41. Morchón R, Carretón E, González-Miguel J, Mellado-Hernández I. Heartworm Disease (Dirofilaria immitis) and Their Vectors in Europe - New Distribution Trends. Front Physiol. 2012;3.

42. Wehmeyer ML, Jaworski L, Jöst H, Șuleșco T, Rauhöft L, Afonso SMM, et al. Host attraction and host feeding patterns indicate generalist feeding of Culex pipiens s.s. and Cx. torrentium. Parasites and Vectors. 2024 Dec 1;17(1):1–12.

43. European Centre for Disease Prevention and Control. Mosquito maps [Internet]. 2024 Jun [cited 2024 Jun 28]. Available from: https://www.ecdc.europa.eu/en/disease-vectors/surveillance-and-disease-data/mosquito-maps

44. Börstler J, Jöst H, Garms R, Krüger A, Tannich E, Becker N, et al. Host-feeding patterns of mosquito species in Germany. Parasit Vectors. 2016;9(1):1–14.

45. Fuehrer HP, Morelli S, Unterköfler MS, Bajer A, Bakran-Lebl K, Dwużnik-Szarek D, et al. Dirofilaria spp. and Angiostrongylus vasorum: Current Risk of Spreading in Central and Northern Europe. Pathogens. 2021 Oct 1;10(10).

46. Pampiglione S, Canestri Trotti G, Rivasi F. Human dirofilariasis due to Dirofilaria (Nochtiella) repens: a review of world literature. Parassitologia. 1995 Dec;37(2–3):149– 93.

47. Pampiglione S, Rivasi F. Human dirofilariasis due to Dirofilaria (Nochtiella) repens: an update of world literature from 1995 to 2000. Parassitologia. 2000 Dec;42(3–4):231–54.

48. Lee ACY, Montgomery SP, Theis JH, Blagburn BL, Eberhard ML. Public health issues concerning the widespread distribution of canine heartworm disease. Trends Parasitol. 2010 Apr 1;26(4):168–73.

49. Huang H, Wang T, Yang G, Zhang Z, Wang C, Yang Z, et al. Molecular characterization and phylogenetic analysis of Dirofilaria immitis of China based on COI and 12S rDNA genes. Vet Parasitol. 2009 Mar 9;160(1–2):175–9.

50. Bazzocchi C, Jamnongluk W, O’Neill SL, Anderson TJC, Genchi C, Bandi C. wsp Gene sequences from the Wolbachia of filarial nematodes. Curr Microbiol. 2000;41(2):96–100.

51. Genchi C, Solari Basano F, Bandi C, Di Sacco B, Venco L, Vezzoni A, et al. Proceedings of Heartworm Sympo-sium ’95. American Heartworm Society, Baravia, IL,. 1995. p. 65–71 Factors influencing the spread of heartworms in Italy Interaction between Dirofilaria immitis and Dirofilaria repens.

52. Genchi C, Mortarino M, Rinaldi L, Cringoli G, Traldi G, Genchi M. Changing climate and changing vector-borne disease distribution: The example of Dirofilaria in Europe. Vet Parasitol. 2011 Mar 22;176(4):295–9.

53. Lee ACY, Atkins CE. Understanding feline heartworm infection: disease, diagnosis, and treatment. Top Companion Anim Med. 2010 Nov;25(4):224–30.

54. Mazzariol S, Cassini R, Voltan L, Aresu L, Frangipane Di Regalbono A. Heartworm (Dirofilaria immitis) infection in a leopard (Panthera pardus pardus) housed in a zoological park in north-eastern Italy. Parasit Vectors. 2010;3(1).

55. Alho AM, Marcelino I, Colella V, Flanagan C, Silva N, Correia JJ, et al. Dirofilaria immitis in pinnipeds and a new host record. Parasit Vectors. 2017 Mar 13;10(1).

56. Molnár V, Pazár P, Rigó D, Máthé D, Fok É, Glávits R, et al. Autochthonous Dirofilaria immitis infection in a ferret with aberrant larval migration in Europe. Journal of Small Animal Practice. 2010 Jul;51(7):393–6.

57. Ruiz de Ybáñez MR, Martínez-Carrasco C, Martínez JJ, Ortiz JM, Attout T, Bain O. Dirofilaria immitis in an African lion (Panthera leo). Veterinary Record. 2006 Feb 18;158(7):240–2.

58. Ionicǎ AM, Matei IA, D’Amico G, Ababii J, Daskalaki AA, Sándor AD, et al. Filarioid infections in wild carnivores: a multispecies survey in Romania. Parasit Vectors. 2017 Jul 13;10(1).

59. Penezić A, Selaković S, Pavlović I, Ćirović D. First findings and prevalence of adult heartworms (Dirofilaria immitis) in wild carnivores from Serbia. Parasitol Res. 2014 Sep 1;113(9):3281–5.

60. Tolnai Z, Széll Z, Sproch Á, Szeredi L, Sréter T. Dirofilaria immitis: An emerging parasite in dogs, red foxes and golden jackals in hungary. Vet Parasitol. 2014 Jul 14;203(3–4):339–42.

61. Potkonjak A, Rojas A, Gutiérrez R, Nachum-Biala Y, Kleinerman G, Savić S, et al. Molecular survey of Dirofilaria species in stray dogs, red foxes and golden jackals from Vojvodina, Serbia. Comp Immunol Microbiol Infect Dis. 2020;68.

62. Ćirović D, Penezić A, Pavlovicv I, Kulišić Z, Ćosić N, Burazerović J, et al. First records of dirofilaria repens in Wild Canids from the region of central Balkan. Acta Vet Hung. 2014;62(4):481–8.

63. Marconcini A, Magi M, Macchioni G, Sassetti M. Filariosis in foxes in Italy. Vet Res Commun. 1996;20(4):316–9.

64. Magi M, Calderini P, Gabrielli S, Dell’Omodarme M, Macchioni F, Prati MC, et al. Vulpes vulpes: A possible wild reservoir for zoonotic filariae. Vector-Borne and Zoonotic Diseases. 2008 Apr 1;8(2):249–52.

65. Gavrilović P, Dobrosavljević I, Vasković N, Todorović I, Živulj A, Kureljušić B, et al. Cardiopulmonary parasitic nematodes of the red fox (Vulpes vulpes) in Serbia. Acta Vet Hung. 2019;67(1):60–9.

66. Gortázar C, Villafuerte R, Lucientes J, Fernández-de-Luco D. Habitat related differences in helminth parasites of red foxes in the Ebro valley. Vet Parasitol. 1998 Dec 15;80(1):75–81.

67. Gortazar C, Castillo JA, Lucientes J, Blanco JC, Arriolabengoa A, Calvete C. Factors affecting Dirofilaria immitis prevalence in red foxes in northeastern Spain. J Wildl Dis. 1994;30(4):545–7.

68. Hodžić A, Alić A, Klebić I, Kadrić M, Brianti E, Duscher GG. Red fox (Vulpes vulpes) as a potential reservoir host of cardiorespiratory parasites in Bosnia and Herzegovina. Vet Parasitol. 2016;223:63–70.

69. Morgan ER, Tomlinson A, Hunter S, Nichols T, Roberts E, Fox MT, et al. Angiostrongylus vasorum and Eucoleus aerophilus in foxes (Vulpes vulpes) in Great Britain. Vet Parasitol. 2008 Jun 14;154(1–2):48–57.

70. IonicǍ AM, Matei IA, D’Amico G, Daskalaki AA, Juránková J, Ionescu DT, et al. Role of golden jackals (Canis aureus) as natural reservoirs of Dirofilaria spp. in Romania. Parasit Vectors. 2016 Apr 28;9(1).

71. Gavrilović P, Marinković D, Todorović I, Gavrilović A. First report of pneumonia caused by Angiostrongylus vasorum in a golden jackal. Acta Parasitol. 2017;62(4):880– 4.

72. Miterpáková M, Hurníková Z, Zaleśny G, Chovancová B. Molecular evidence for the presence of Dirofilaria repens in beech marten (Martes foina) from Slovakia. Vet Parasitol. 2013 Sep 23;196(3–4):544–6.

73. Gavrilović P, Blitva-Robertson G, Özvegy J, Kiskároly F, Becskei Z. Case report of dirofilariasis in gray wolf in Serbia. Acta Parasitol. 2015 Jan 1;60(1):175–8.

74. Hamel D, Silaghi C, Lescai D, Pfister K. Epidemiological aspects on vector-borne infections in stray and pet dogs from Romania and Hungary with focus on Babesia spp. Parasitol Res. 2012 Apr;110(4):1537–45.

75. Ciucă L, Musella V, Miron LD, Maurelli MP, Cringoli G, Bosco A, et al. Geographic distribution of canine heartworm (Dirofilaria immitis) infection in stray dogs of eastern Romania. Geospat Health. 2016;11(3):318–23.

76. Stoyanova H, Carretón E, Montoya-Alonso JA. Stray Dogs of Sofia (Bulgaria) Could be an Important Reservoir of Heartworm (Dirofilaria Immitis). Helminthologia. 2019 Nov 5;56(4):329–33.

77. Fortin JF, Slocombe JOD. Temperature requirements for the development of Dirofilaria immitis in Aedes triseriatus and Ae. vexans. Mosq News. 1981;41(4):625–33.

78. Cancrini G, Pietrobelli M, Frangipane di Regalbono AF, Tampieri MP, della Torre A. Development of Dirofilaria and Setaria nematodes in Aedes albopictus. Parassitologia. 1995 Dec;37(2–3):141–5.

79. Miranda MÁ, Barceló C, Arnoldi D, Augsten X, Bakran-Lebl K, Balatsos G, et al. AIMSurv: First pan-European harmonized surveillance of Aedes invasive mosquito species of relevance for human vector-borne diseases. GigaByte (Hong Kong, China). 2022 May 31;2022:1–11.

80. Benedict MQ, Levine RS, Hawley WA, Lounibos LP. Spread of the tiger: global risk of invasion by the mosquito Aedes albopictus. Vector Borne Zoonotic Dis. 2007 Mar;7(1):76–85.

81. Oliveira S, Rocha J, Sousa CA, Capinha C. Wide and increasing suitability for Aedes albopictus in Europe is congruent across distribution models. Sci Rep. 2021 Dec 1;11(1).

82. Companion Animal Parasite Council | Heartworm [Internet]. [cited 2023 Jul 29]. Available from: https://capcvet.org/guidelines/heartworm/

